# Ontology-based validation and identification of regulatory phenotypes

**DOI:** 10.1101/256529

**Authors:** Maxat Kulmanov, Paul N Schofield, Georgios V Gkoutos, Robert Hoehndorf

**Affiliations:** Computer, Electrical and Mathematical Sciences and Engineering Division, Computational Bioscience Research Center, King Abdullah University of Science and Technology, Thuwal 23955, Kingdom of Saudi Arabia; Department of Physiology, Development & Neuroscience, University of Cambridge, Downing Street, CB2 3EG, Cambridge, United Kingdom; College of Medical and Dental Sciences, Institute of Cancer and Genomic Sciences, Centre for Computational Biology, University of Birmingham, B15 2TT, Birmingham, United Kingdom; Institute of Translational Medicine, University Hospitals Birmingham, NHS Foundation Trust, B15 2TT, Birmingham, United Kingdom; Institute of Biological, Environmental and Rural Sciences, Aberystwyth University, SY23 2AX, Aberystwyth, United Kingdom

## Abstract

**Motivation:** Function annotations of gene products, and phenotype annotations of genotypes, provide valuable information about molecular mechanisms that can be utilized by computational methods to identify functional and phenotypic relatedness, improve our understanding of disease and pathobiology, and lead to discovery of drug targets. Identifying functions and phenotypes commonly requires experiments which are time-consuming and expensive to carry out; creating the annotations additionally requires a curator to make an assertion based on reported evidence. Support to validate the mutual consistency of functional and phenotype annotations as well as a computational method to predict phenotypes from function annotations, would greatly improve the utility of function annotations.

**Results:** We developed a novel ontology-based method to validate the mutual consistency of function and phenotype annotations. We apply our method to mouse and human annotations, and identify several inconsistencies that can be resolved to improve overall annotation quality. Our method can also be applied to the rule-based prediction of phenotypes from functions. We show that the predicted phenotypes can be utilized for identification of protein-protein interactions and gene-disease associations. Based on experimental functional annotations, we predict phenotypes for 1,986 genes in mouse and 7,301 genes in human for which no experimental phenotypes have yet been determined.

**Availability:** https://github.com/bio-ontology-research-group/phenogocon

## 1 Introduction

Although several definitions of what constitutes a phenotype have been proposed over time, a phenotype can be operationally defined as an observable characteristic of an organism arising from interactions between the organism’s genotype and the environment (Johannsen, 1909, 1911). Understanding the molecular and functional basis of phenotypes is an important factor in our understanding of disease mechanisms.

Abnormal phenotypes associated with loss of gene function provide valuable information for a variety of computational methods, such as identification of gene-disease associations (Hirschhorn *et al.*, 2002), protein-protein interactions (Kahanda *et al.*, 2015; Hu *et al.*, 2011), disease causative variant prioritization (Boudellioua *et al.*, 2017), finding orthologous genes (Hoehndorf *et al.*, 2011), and drug discovery (Moffat *et al.*, 2014) and repurposing (Hoehndorf *et al.*, 2014). Identifying which phenotypes a gene may be associated with is challenging; even in the case of a complete loss of function of a gene, phenotypes may be highly variable (de Angelis *et al.*, 2015).

Several consortia and research initiatives aim to systematically catalog the phenotypes associated with loss of function mutations in model organisms (Ring *et al.*, 2015), and the experimental results produced by these initiatives provide valuable information for understanding gene function (Ring *et al.*, 2015)or their role in disease (Meehan *et al.*, 2017). In addition to high-throughput phenotyping, there are also ongoing efforts to identify genotype–phenotype relations from literature (Smith and Eppig, 2015), and to record phenotypes observed in a clinical setting which are associated with particular genotypes (Landrum *et al.*, 2013).

There are several computational methods available for predicting the functions of proteins (Kulmanov *et al.*, 2017; Cozzetto *et al.*, 2016; Gong *et al.*, 2016). Computational methods for function prediction have improved in predictive performance and, subsequently, in their utility, over recent years (Radivojac *et al.*, 2013). Consequently, it is a reasonable question to ask whether the same or similar approaches may also work for phenotypes, i.e., whether we can build efficient methods to predict phenotypes from genotypes, and whether these methods can provide information that may be of clinical utility. While methods for protein function prediction are maturing, computational methods to predict phenotypes are still in their infancy.

There are many challenges in predicting phenotypes, both biologically and computationally. From a biological perspective, predicting the phenotypes that arise from a particular genotype is challenging due to the complex molecular and physiological interactions that give rise to phenotypes, open-ended environmental influences and determinants of phenotypes, incomplete penetrance and resilience of organisms to certain phenotypic manifestations, epigenetic regulation not detectable on the level of a genotype, and many other factors contributing to the variability and heterogeneity of phenotypes. The impact of pleiotropy and genetic background were themselves instrumental in motivating the very large scale knockout mouse project (IKMC), precisely because of the problems intrinsic to predicting phenotype from genotype (Tyler *et al.*, 2016; Austin *et al.*, 2004).

From a computational perspective, there are also several additional challenges. First, there is a substantial lack of potential training data that limits the application of machine learning approaches. The high variability in phenotypes and their descriptions (Gkoutos *et al.*, 2005) makes it challenging to identify whether genotypes are involved in identical or similar phenotypes. There is also a lack of computationally represented background knowledge necessary to determine the relationship between phenotypes and their physiological and patho-physiological basis; in particular, there is no computationally accessible, qualitative representation of physiological interactions in mammals. Furthermore, representation of environmental influences is challenging, partly due to their heterogeneity, but also failure to capture environmental parameters in many phenotyping studies (Beckers *et al.*, 2009; Schofield *et al.*, 2016).

The premise underlying comprehensive phenotyping studies is that, uniquely, the phenotype of an organism lacking a functioning copy of a given gene provides definitive information on gene function; the primary goal of functional genomics. Here, we investigate the relationship between Gene Ontology (GO) (Ashburner *et al.*, 2000) functions that are associated with gene products, and phenotypes associated with a loss of function in these gene products (either through targeted or random mutation, epigenetic modification or pharmaceutical effects). Our aim is to identify how much information functions of gene products carry about the phenotypes in which these gene products are involved. Specifically, we test the hypothesis that a loss of a regulatory function (i.e., the up-or down-regulation of some other process) will result in a regulatory phenotype. For example, if a protein is (unconditionally) involved in a *positive regulation of B cell apoptosis*, then a loss of function in that protein should lead to a phenotype in which the rate of *B cell apoptosis* is decreased. We first formalize our assumptions in rules that relate axioms in the Web Ontology Language (OWL) (Grau *et al.*, 2008). We then test how many function – phenotype pairs in the laboratory mouse (*Mus musculus*) and the human (*Homo sapiens*) satisfy these rules, how many annotations are consistent with our hypothesis, and how many annotations are not consistent with out hypothesis. We investigate some of the inconsistent pairs we identify, and characterize the reasons for the inconsistency; we find that they can be a result of incomplete or under-specified contextualization of function or phenotype annotations (such as by cell type), conflicting annotation derived from literature, or a consequence of inference over the ontology structure.

After validating and characterizing possible inconsistent annotations, we apply our hypothesis predictively and predict regulatory phenotypes associated with loss of function mutations in 11,987 gene products in the mouse and 15,680 in the human. We validate our predictions by predicting gene-disease associations and protein-protein interactions from phenotypes and demonstrate that our rules result in predictions that are predictive of known associations.

## 2 Methods

### 2.1 Data sources

We use functional and phenotypic annotations for mouse and human. We downloaded Gene Ontology (GO) (Ashburner *et al.*, 2000) annotations from http://geneontology.org/ on December 5th 2017. The file contains 439,128 distinct annotations to 19,452 human gene products, and 376,532 distinct annotations to 24,526 mouse gene products. We use the phenotype annotations for mouse downloaded from the Mouse Genome Informatics (MGI) (Smith and Eppig, 2015) database (http://www.informatics.jax.org/downloads/reports/index.html) on December 5th 2017. We use the *MGI_Gene_Pheno.rpt* file which contains phenotypes for non-conditional loss of function mutations in single genes; the file contains phenotypes for 11,887 mouse genes and 206,272 distinct associations between a gene and a Mammalian Phenotype Ontology (MP) (Smith and Eppig, 2015) class. For human, we downloaded annotations provided by the Human Phenotype Ontology (HPO) database (Robinson *et al.*, 2008) on December 5th 2017. We use the file containing phenotypes from “all sources”and “all frequencies”; the file contains phenotype associations for 3,682 human genes and 120,289 distinct associations between human genes and HPO classes.

For reasoning and processing formal definitions of phenotypes, we use the multi-species integrated PhenomeNET ontology (Hoehndorf *et al.*, 2011; Rodríguez-García *et al.*, 2017). We downloaded the latest version of the PhenomeNET ontology from the AberOWL (Hoehndorf *et al.*, 2015) ontology repository http://aber-owl.net/ontology/PhenomeNET/. We also downloaded the GO in its OWL format, released on December 2nd 2017, from the AberOWL ontology repository.

### 2.2 Filtering GO annotations

To obtain only experimental GO annotations, we filtered all GO annotations by their evidence codes so that we only retain annotations with an experimental evidence. Specifically, we only keep annotations with evidence codes *EXP*, *IDA*, *IPI*, *IMP*, *IGI*, *IEP*, *TAS*, and *IC*. We removed all annotations which are negated (i.e., using a NOT qualifier); we also excluded all annotations that are context specific, i.e., which are explicitly conditional on a particular environment or other restrictions (such as occurring only in particular cell types, or tissues, or during certain developmental stages).

After filtering all annotations, our GO annotation set contains 100,336 annotations to 11,987 mouse gene products and 295,357 annotations to 15,680 human gene products. We mapped all protein identifiers to MGI identifiers for mouse proteins, and to HUGO (Yates *et al.*, 2017) standard human gene names.

### 2.3 Gene-Disease associations

Our phenotype predictions are evaluated by assessing how well they permit recapitulation of known gene-disease relationships based on phenotype similarity. Specifically, we predict association between mouse and human genes and Mendelian diseases from the Online Mendelian Inheritance in Man (OMIM) database (Amberger *et al.*, 2011); we use the associations between mouse genes and OMIM diseases from the MGI database, and the associations provided by the HPO team for human genes and OMIM diseases, downloaded on 27 January 2018.

We downloaded phenotypes associated with diseases from the HPO database *phenotype_annotation* file provided by HPO team from http://compbio.charite.de/jenkins/job/hpo.annotations.monthly/lastStableBuild/ on January 27th 2017. For human genes, we used phenotype annotations of homologous genes in the mouse from the MGI database.

### 2.4 Protein-Protein Interaction

For further validation of our predictions, we use protein-protein interactions provided by the STRING Database (Szklarczyk *et al.*, 2015). We downloaded all mouse and human interactions from STRING version 10.5 and filtered the interactions by a confidence score higher or equal to 300. We use the protein.aliases file provided by the STRING database to map STRING protein identifiers to MGI identifiers (for mouse genes and proteins) and HUGO gene names (for human genes and proteins).

### 2.5 Computing semantic similarity

We measure the similarity between sets of MP and HPO classes by computing Resnik’s pairwise similarity measure using the PhenomeNET Ontology (Hoehndorf *et al.*, 2011), and using the Best-Match-Average (BMA) (Pesquita *et al.*, 2009) strategy to combine pairwise similarities into a single similarity score between two sets of annotations. We use the normalized similarity value as a prediction score for PPI or gene-disease association and compute the area under the receiver operating characteristic (ROC) curve (Fawcett, 2006) as a quantitative measure of predictive performance.

Resnik’s similarity measure uses the information content (IC). IC is computed as the probability of occurrence of a class in annotations:

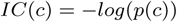

The similarity value between two classes is the IC of the most informative common ancestor (MICA), i.e.:

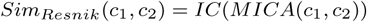

For two sets of classes we compute the similarity value between each pair and use the BMA combination strategy:

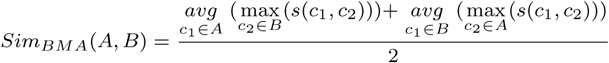

where *s*(*x*, *y*) = *Sim*_*Resnik*_(*x*, *y*).

### 2.6 Predicting protein functions with DeepGO

In order to evaluate our method for predicting phenotypes from functions for gene products without experimental annotations, we predicted GO function annotations using the DeepGO function prediction system (Kulmanov *et al.*, 2017). We downloaded SwissProt reviewed human and mouse protein sequences from the UniProt database (The UniProt Consortium, 2017) on 28 January 2018.

Initially, our dataset had 16,950 mouse and 20,244 human proteins. To meet the DeepGO requirements and limitations, we filtered this set of proteins and removed all sequences with ambiguous amino acid symbols (i.e., B, O, J, U, X, Z); we also removed all proteins with more than 1002 amino acids. After filtering, we retained 14,916 mouse and 17,837 human proteins for which we could predict functions using DeepGO. We mapped UniProt identifiers to MGI identifiers and HUGO gene names.

### 2.7 Implementation

We implemented our approach using the OWL API (Horridge and Bechhofer, 2011) version 4.1.0 and used the Similarity Measures Library (Harispe *et al.*, 2014) for measuring semantic similarities. The source code, documentation, and data files are freely available at https://github.com/bio-ontology-research-group/phenogocon.

## 3 Results

### 3.1 The correspondence between regulation and regulatory phenotypes

Our main hypothesis is that there should be a close relationship between some functions to which gene products are annotated and some phenotypes. In particular, if a gene product is involved in the up-or down-regulation of a process *P*, then a loss-of-function of that gene product (introduced, for example, through a pathogenic variant, a targeted mutation, or an epigenetic interference) will usually lead to a phenotype in which the rate or intensity of *P* is decreased or increased.

Specifically, we assume that, if a phenotype is defined as a change of some biological process (such as an increased or decreased rate or turnover of the process), then we can annotate the gene products which negatively or positively regulate or contribute to *P* biological process with the given phenotype. For example, when a protein that is normally involved in *positive regulation of B cell apoptotic process* (`GO:0002904`) is inhibited (for example through a genetic mutation, or through a small molecule which inhibits the protein), we would expect that the rate with which processes of the type *B cell apoptotic process* (`B cell apoptotic process`) occur to decrease.

We formalize this hypothesis in the form of rules that assign a new annotation to a protein with a particular function annotation. Let *X* be a protein involved in (i.e., annotated with) the function *P*. We then implement our hypothesis through the following three rules:

- **Increase Function – Decreased Phenotype**: If `P SubClassOf ‘positively regulates’ some P2`, then a loss of function of *X* results in the phenotype `‘phenotype of’ some (P2 and ‘has quality’ some ‘decreased quality’)`.
- **Decreased Function – Increased Phenotype**: If `P SubClassOf ‘negatively regulates’ some P2`, then a loss of function of *X* results in the phenotype `‘phenotype of’ some (P2 and ‘has quality’ some ‘increased quality’)`.
- **Abnormal Function – Abnormal Phenotype**: A loss of function of *X* results in the phenotype `‘phenotype of’ some (P and ‘has quality’ some (‘has modifier’ some abnormal))`

While the first two rules directly implement our hypothesis, the third rule establishes a correspondence between a loss of GO function and the resulting phenotype; it is, in a sense, more general than the previous two rules which establish a correspondence between regulatory functions and phenotypes. The inverse of the abnormality rule has already been used to predict GO functions from phenotypes (Hoehndorf *et al.*, 2013).

To determine whether a pair of classes in GO and a phenotype ontology class match our hypothesis and subsequent rules, we use the formal definitions and axioms that constrain the GO classes and the classes in phenotype ontologies. Many classes in phenotype ontologies are formally defined using definition patterns based on the Entity–Quality (EQ) method (Gkoutos *et al.*, 2005; Mungall, 2009; Gkoutos *et al.*, 2017). In the EQ method, phenotypes are decomposed into an entity – either an anatomical entity or a biological process or function – and a quality. We identify the GO class underlying each phenotype in MP and HPO based on these EQ-based definition patterns, and we also identify for each phenotype the direction (i.e., *increased or decreased*) in which the process or function is modified. As a result, we obtain, for each phenotype in HPO or MP that is based on an abnormal function or process, a pair of a GO class and a direction.

For example, the class *Increased thymocyte apoptosis* (`MP:0009541`) is defined using the Entity *Thymocyte apoptotic process* (`GO:0070242`) and the Quality *Increased rate* (`PATO:0000912`); the Quality is further constrained by adding the *Abnormal* (`PATO:0000460`) quality (in order to distinguish the abnormal phenotype from a physiological increase in thymocyte apoptotic rate).

In total, there are 287 phenotype classes in HPO based on GO processes or functions, of which 17 are increased in rate, 54 are decreased in rate, and 216 are abnormalities of a process or function. In MP, 1,543 classes are based on GO processes or functions, of which 272 classes are increased in rate, 342 classes are decreased in rate, and 929 classes are abnormalities of a process or function.

As next step in our workflow, we identify all GO processes that up-or down-regulate other processes. For this purpose, we use the Elk OWL reasoner (Kazakov *et al.*, 2012) to query GO for all equivalent classes of ‘`Biological regulation’ and ‘positively regulates’ some X` and `‘Biological regulation’ and ‘negatively regulates’ some X` for all classes X. In total, we identify 3,013 processes that positively regulate another biological process, and 3,043 processes that negatively regulate another biological process.

In total, we identify 1, 570 correspondence rules between GO and phenotype classes of which 1, 328 classes are from MP and 242 classes are from HPO. The complete set of correspondences between a GO class and phenotype class is available on our project website. We use the correspondences between regulatory phenotypes and GO functions in two ways: first, we evaluate how many annotations are inconsistent with these rules, and determine why they are inconsistent; second, we use these rules to predict phenotypes from GO functions.

### 3.2 Determining consistency between function annotations and phenotype annotations

We define a consistency between regulation functions and regulatory phenotypes as annotations that do not contradict our rules, i.e., when either no evidence about a phenotype (or function) is provided in the annotations, or when they correspond to our rules. An inconsistent pair of annotations is a pair of function and phenotype annotations which contradict our rules (i.e., the function annotation is to the up-or down-regulation of a process and the phenotype of the loss of function is an increased or decreased rate of that process). We generated 423 GO–phenotype pairs that represent an inconsistency; of these 423 pairs, 398 pairs are GO–MP classes and 25 pairs are GO–HPO classes.

We determine whether the function and phenotype annotations in the Mouse Genome Informatics (MGI) (Smith and Eppig, 2015) model organism database are consistent with our hypothesis, and whether the function annotations for human proteins provided by UniProt (The UniProt Consortium, 2017) and the phenotypes associated with these proteins provided by the HPO database (Robinson *et al.*, 2008; Köhler *et al.*, 2017) are consistent. In the first instance, and to identify only unambiguously matching pairs, we ignore inferences over the ontology and consider only exactly matching phenotypes, i.e., only the annotations in which the direct annotation to the phenotype matches our rule. We find 105 function– phenotype annotation pairs for mouse and one annotation for human which are inconsistent according to our set of inconsistent pairs.

We manually analyzed some of the annotations we tagged as inconsistent with our rules. In many cases, inconsistency with our rules may arise from conflicting GO or phenotype annotations. For example, folliculin interacting protein 1 (*Fnip1*, `MGI:2444668`) is annotated with the GO function *Positive regulation of B cell apoptotic process* (`GO:0002904`), and the loss of function of *Fnip1* is annotated with the phenotype *increased B cell apoptosis* (`MP:0008782`). Using our rule (Increased Function – Decreased Phenotype), we flagged this pair of annotations as inconsistent. Both annotations are asserted based on evidence from the same publication (Park *et al.*, 2012), which reports a negative regulatory role for *Fnip1* in B cell apoptosis and uses as experimental evidence that B cell apoptosis is increased in response to metabolic stress in mice lacking *Fnip1* function. The reports in the paper, together with our rule-based identification of the possible inconsistency, indicates that the GO annotation of *Fnip1 to Positive regulation of B cell apoptotic process* may not be correct and should be replaced by an annotation to *Negative regulation of B cell apoptotic process*.

Another example involved glypican 3 (*Gpc3*, `MGI:104903`), which is annotated with the function *Negative regulation of growth* and the phenotype *Postnatal growth retardation*. Here, the asserted annotation to postnatal growth retardation is based on Chiao *et al.* (2002). The postnatal growth catch-down and catch-up seen in homozygote nulls was subject to extensive analysis in the paper and the authors conclude that the normal, growth suppressing, function of *Gpc3* is restricted to the embryonic period. The knockout phenotype should therefore have been annotated as *Increased embryo size*, not *Postnatal growth retardation* as the closest description to the phenotype described in the paper.

The complexity of phenotypic annotations is well demonstrated by the inconsistency we detect for an annotation of the CD28 cell surface receptor. Annotated in GO to *Positive regulation of T cell proliferation*, the knockout strain phenotype is annotated in MGI to *Increased T cell proliferation* (Bour-Jordan *et al.*, 2004). Regulatory T cells (Tregs; CD4+CD25+) depend on CD28 for activation and proliferation. Effector T cells are suppressed in non-obese diabetic (NOD) mice by active Tregs. In the absence of CD28, Tregs do not proliferate, thereby permitting effector cells to proliferate. This proliferation of effector T cells is reported in the manuscript on which the phenotype annotation is based, and leads to the phenotype annotation of the knockout. Formally this is accurate, but the phenotype reported is dependent on the function of a cell type whose own function is affected by the loss of CD28 in a different cell. This “russian doll”effect is likely to be a significant confounder in relating phenotype to function, particularly at a high level of phenotypic granularity.

We also experimented with extending the scope of our method and included inferred phenotype annotations (we consider a phenotype annotation to phenotype class *C* as inferred if and only if the annotation is made to a subclass of *C* in the phenotype ontology). This allows us to identify significantly more potentially inconsistent function – phenotype pairs. We find, for example, the inconsistent annotation pair in BCL2-associated athanogene 6 (*BAG6*) between the GO process *Negative regulation of apoptotic process* (`GO:0043066`) and *Decreased apoptosis* (`MP:0006043`). However, the directly asserted annotation of *BAG6* is to *Decreased susceptibility to neuronal excitotoxicity* (`MP:0008236`), a subclass of *Decreased apoptosis* in MP. While a direct annotation to *Decreased apoptosis* would likely have implied that apoptotic processes are, in general, decreased in rate, an annotation to *Decreased susceptibility to neuronal excitotoxicity* does not have the same implications: apoptotic processes occurring in neurons under certain conditions are decreased in rate, but most apoptotic processes are unaffected. Due to these implications, we do not apply our rules to phenotypes that are inferred over a phenotype ontology.

### 3.3 Predicting phenotypes from functions

We can also use our rules to predict phenotypes from function annotations. In this case, we take function annotations of a gene product as input, and predict a phenotype that satisfies the definition in our rules. Not all function annotations readily imply a phenotype; therefore, we cannot generate phenotype annotations for all proteins. We generated 78,298 phenotype annotations for 10,041 human genes, and 61,875 phenotype annotations for 7,314 mouse genes. Of the generated annotations, 116 human gene annotations and 3,170 mouse gene annotations are already present in our data while the remaining predictions are novel. Notably, we predict phenotypes for 1,986 genes that have no phenotypes at all in the mouse, and for 7,301 genes without any phenotype annotations in the human. Table 1 summarizes our findings.

**Table 1.**
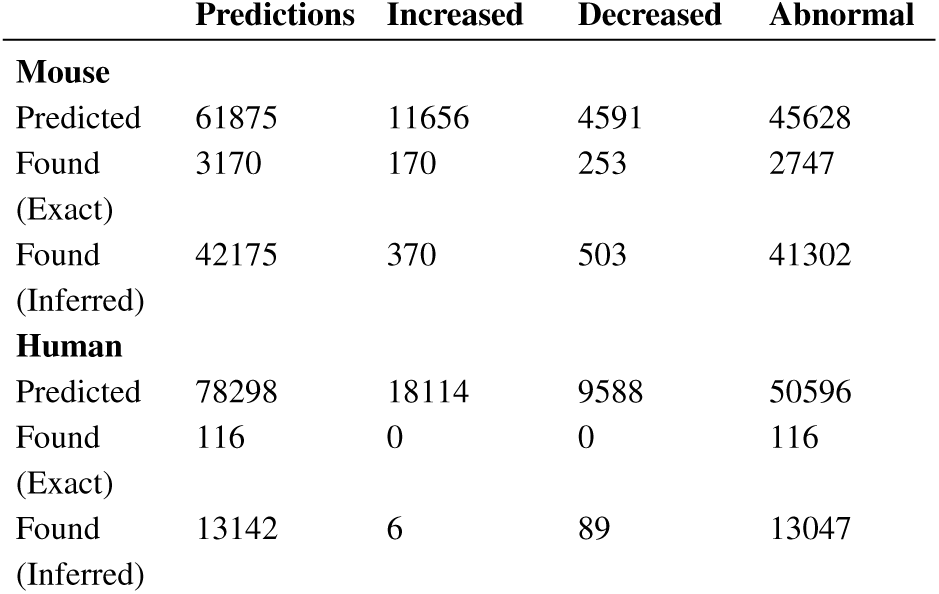
Number of predicted annotations using exact matching rules and rules inferred with ontology structure, and the number of annotations that are already asserted. For inferred matches we assume that genotypes are annotated to all superclasses of their annotated classes and propagate both functional and phenotypic annotations. For example, if a genotype has the phenotype Increased B cell apoptosis and application of our rule predicts increased apoptosis, we will also consider this as a match.

Phenotype annotations have many applications; in particular, it is accepted that phenotypes reflect underlying physiological interactions and networks (Costanzo *et al.*, 2016) and phenotype annotations are widely used to investigate the molecular basis of diseases (Köhler *et al.*, 2009; Singleton *et al.*, 2014). To validate our phenotype predictions, we performed two experiments. First, we apply a measure of semantic similarity to compute the pairwise similarity between phenotypes associated with genes and diseases, and we evaluate how well this similarity recovers known gene–disease associations. Second, we compute the pairwise similarity between phenotypes associated with genes, and we use the gene–gene phenotypic similarity to predict interactions between the genes (combining different interaction types aggregated in the STRING database (Szklarczyk *et al.*, 2015), including genetic interactions and protein–protein interactions).

To predict gene–disease associations, we perform two experiments. First, we recover mouse models of human diseases as characterized in the MGI database, and second, we predict genes associated with diseases in the HPO database (Köhler *et al.*, 2016). We evaluate our performance using a receiver operating characteristic (ROC) curve (Fawcett, 2006). A ROC curve is a plot of a classifiers true positive rate as a function of the false positive rate, and the area under the ROC curve (ROCAUC) is a quantitative measure of a classifier’s performance (Fawcett, 2006). The ROC curves obtained for predicting mouse models of human disease are shown in Figure 1, and the ROC curves for predicting gene–disease associations in humans in Figure 2. We find that when we use only our predicted sets of phenotypes, we can predict gene–disease associations with a ROCAUC of 0.65 (to identify mouse models of human disease) or 0.63 (to identify human gene–disease associations), which is a weak but positive predictive signal. When we merge our predictions and the original phenotype annotations, predictive performance slightly drops in comparison to using only original annotations.

**Fig. 1:**
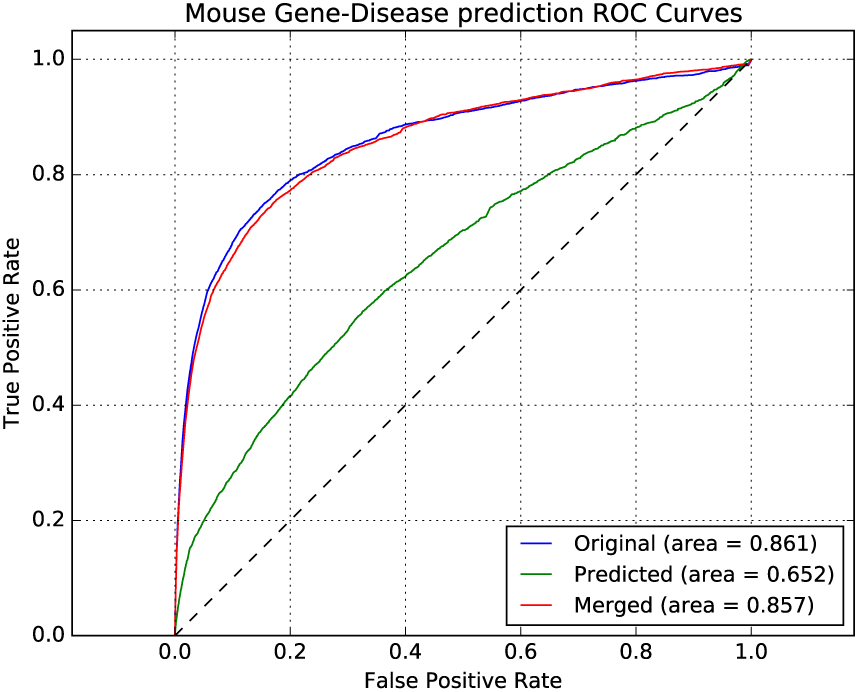
ROC Curves for predicting Gene-Disease associations for mouse genes. *Original* uses asserted phenotype annotations, *Predicted* uses only predicted phenotypes, and *Merged* combine asserted and predicted phenotypes.

**Fig. 2:**
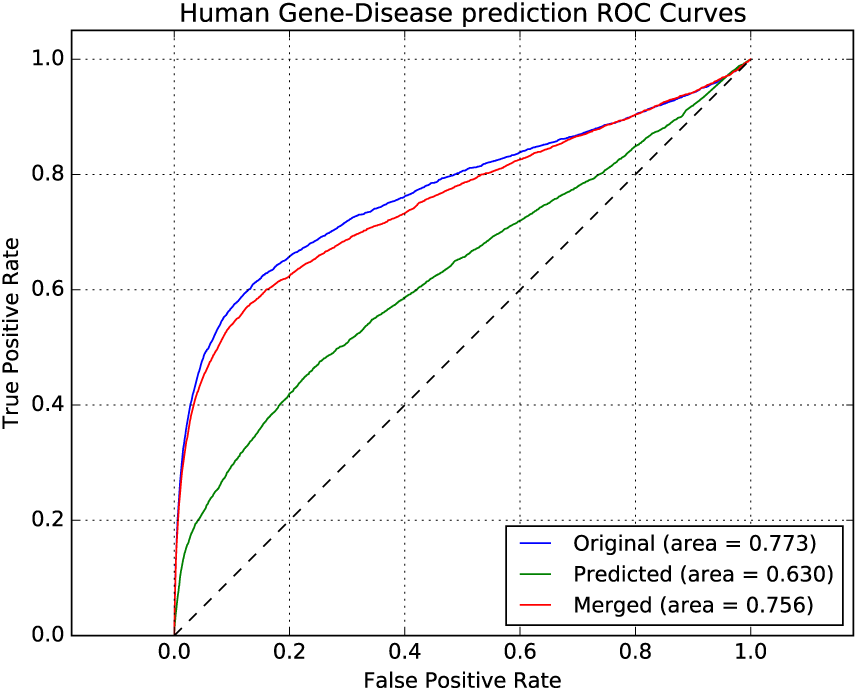
ROC Curves for predicting Gene-Disease associations for human genes. *Original* uses asserted phenotype annotations, *Predicted* uses only predicted phenotypes, and *Merged* combine asserted and predicted phenotypes.

In addition to predicting gene–disease associations, we compute the pairwise phenotype-similarity between genes to predict interactions between genes and proteins. We again use a ROC curve to evaluate the predictive performance. Notably, the performance for predicting interactions improved even over the performance achieved with the original annotations when using the phenotypes generated by our method. Performance further improved when merging original phenotypes and the phenotypes we predict, demonstrating that there is significant complimentary information in both (see Table 2, and Figures 3 and 4).

**Table 2.** Summary of evaluation of prediction phenotypes for mouse and human. Original uses asserted phenotype annotations, Predicted uses only predicted phenotypes, and Merged combine asserted and predicted phenotypes.

| Method | AUC<br>(original) | AUC<br>(predicted) | AUC<br>(merged) |
| --- | --- | --- | --- |
| **Mouse** |  |  |  |
| Gene-Disease association | 0.861 | 0.652 | 0.857 |
| PPI with experimental GO annotations | 0.667 | 0.672 | 0.705 |
| PPI with DeepGO annotations | 0.667 | 0.696 | 0.694 |
| **Human** |  |  |  |
| Gene-Disease association | 0.773 | 0.630 | 0.756 |
| PPI with experimental GO annotations | 0.616 | 0.749 | 0.902 |
| PPI with DeepGO annotations | 0.616 | 0.741 | 0.928 |

**Fig. 3:**
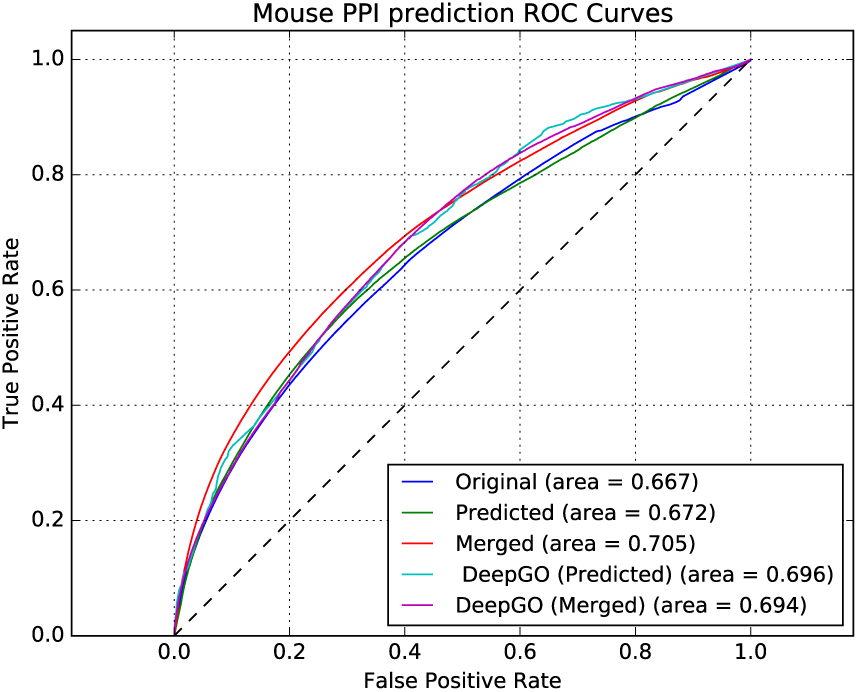
Predicting Protein-Protein interactions using predicted phenotypes for mouse. *Original* uses asserted phenotype annotations, *Predicted* uses only predicted phenotypes, and *Merged* combine asserted and predicted phenotypes. *DeepGO* (*Predicted*) uses only predicted phenotypes based on DeepGO’s predicted GO function annotations, and *DeepGO* (*Merged*) combines them with asserted phenotype annotations.

**Fig. 4:**
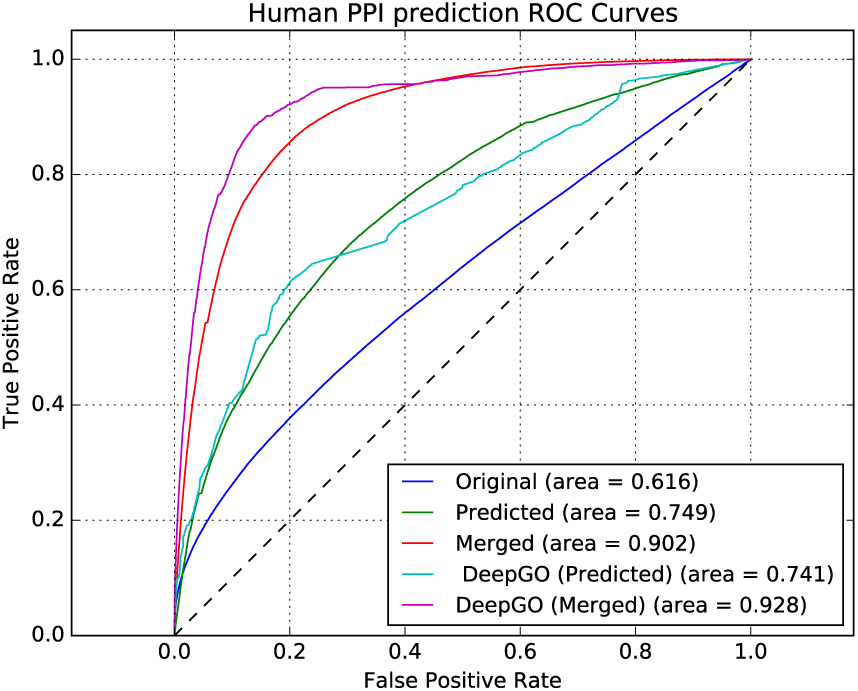
Predicting Protein-Protein interactions using predicted phenotypes for human. *Original* uses asserted phenotype annotations, *Predicted* uses only predicted phenotypes, and *Merged* combine asserted and predicted phenotypes. *DeepGO* (*Predicted*) uses only predicted phenotypes based on DeepGO’s predicted GO function annotations, and *DeepGO* (*Merged*) combines them with asserted phenotype annotations.

### 3.4 Predicting functions, predicting phenotypes

Our method mainly relies on functional annotations of gene products. However, not all genes and gene products have experimental functional annotations. Furthermore, the manual annotations are often derived from mutant phenotypes, thereby limiting the scope of our approach. However, with the recent advances in methods for computational prediction of protein functions (Kulmanov *et al.*, 2017; Cozzetto *et al.*, 2016; Gong *et al.*, 2016; Radivojac *et al.*, 2013), we can experiment with a two-step process: first, we predict GO functions for proteins, and, second, we predict phenotypes arising from a loss of function in the protein using our rules.

We recently developed DeepGO (Kulmanov *et al.*, 2017), a computational method for function prediction which uses a deep neural network algorithm to predict functions from protein sequence and (when available) a cross-species interaction network. Using DeepGO, we can predict functions for gene products with known amino-acid sequences. From the predicted function, we can predict phenotypes using our rules.

The DeepGO model can only predict annotations to 932 distinct biological process classes in GO (Kulmanov *et al.*, 2017). Of the 932 classes that DeepGO can predict, 443 classes are covered by our rules, and 28 classes are negative regulations and 55 classes are positive regulations.

We used DeepGO to predict at least one function for 14,916 mouse and 17,837 human proteins, and based on them, we generated phenotype annotations for 13,225 mouse and 14,187 human genes. 6,033 mouse genes and 11,570 human genes for which we predicted phenotypes do not currently have any experimental phenotype annotations.

We evaluated our predictions using interactions from the STRING database, similarly to our evaluation of phenotypes predicted from experimental GO annotations. Figures 3 and 4 show the performance of predicting interactions in mouse and human, respectively. We find that predicting phenotypes based on DeepGO’s predicted functions allows us to further improve our ability to predict interactions in humans. For the mouse, however, the performance of predicting interactions using phenotypes generated from DeepGO’s predicted functions is slightly lower than predictions based on experimental GO annotations. Table 2 provides a summary of the results.

## 4 Discussion

### 4.1 Rules and statistical approaches for predicting phenotypes

Accurate prediction of the phenotypes of an organism from its genotype, and possibly some environmental features, is probably unachievable in the foreseeable future. However, *some* phenotypes are sufficiently fundamental that they can be predicted reliably given some basic knowledge about a gene and the gene products it encodes. We identify three rules that establish a correspondence between functions of gene products and the phenotypes that a loss of function in these gene products would entail. We believe these rules to be sufficiently robust to hold universally, almost as a consequence of the definition of the corresponding phenotypes. The main limitation in applying our rules predictively is the precision with which function annotations are contextualized, i.e., how universally a function annotation without any context constraints should be interpreted.

There are likely more rules that can be used to reliably predict phenotypes from functions; some may be as simple as the rules we propose, while others may require complex combinations of functions, and additional constraints, to be applied. Rule mining techniques (Bodenreider *et al.*, 2005), in particular those that can utilize axioms and rules in OWL (Lehmann, 2009), could identify more rules of varying strength and may provide an opportunity to further extend our approach.

We demonstrated that we could not only apply our rules to experimentally determined function predictions, but we were also able to use a function prediction method to predict GO functions, then apply our rules and predict phenotypes. While this approach already yields phenotypes that are useful in computational methods (such as similarity-based prediction of protein-protein interactions), some technical modifications could further improve the accuracy and coverage of predicting phenotypes. A main limitation is that both parts of the method are trained and generated separately; an end-to-end learning approach in which phenotypes are predicted directly (and in which the DeepGO model – or another function prediction method – is used as intermediate, pre-trained part) may significantly improve the performance.

### 4.2 What do phenotype annotations mean?

Our method can be used both to identify possibly conflicting annotations as well as to suggest phenotypes that may arise from a particular genotype.

One observation from our experiments is that the meaning of the annotation relation can be different depending on whether the annotation is *asserted* or *inferred* using the ontology structure. Specifically, there seems to be a difference between annotations to a phenotype such as *Increased apoptosis*, depending on whether the annotation is inferred from the ontology hierarchy (as in the case of an annotation to *Increased B cell apoptosis*), or asserted. If the annotation is asserted at the level of the least specific class, we would usually expect *all* types of apoptosis processes in the organism to be increased in rate, including apoptosis of B cells and other specific cell types. However, if the annotation is to a more specific class (such as *increased B cell apoptosis* from which an annotation to *Increased apoptosis* can be inferred, this no longer holds true.

We can use OWL to provide the outlines of a data model in which these considerations are made explicit. Let us assume that *X* is annotated with the phenotype *P*, and, without loss of generality, that *P* is defined as an increased rate of process *F*. There are multiple different options for formalizing the meaning of this annotation. The “weakest”form of interpretation (i.e., the form from which the least amount of information can be derived) would be that an organism with *X* (e.g., an organism with a loss of function mutation in *X*) would have a part in which at least one process of type *F* can be observed to be increased in rate; formally, the organism with *X* would be a subclass of `has-part some ((inverse occurs-in) some (F and has-quality some ‘increased rate’))).` A stronger interpretation could be that *all* processes of type *F* occurring in an organism with *X* would be increased in rate. In this case, processes of type *F* that occur in an organism with *X* would be come a subclass of things with increased rate, i.e, `(F and occurs-in some X) SubClassOf: has-quality some ‘increased rate’.`

From the first interpretation and its formal representation, we cannot conclude that processes of type *F* will always, or usually, be increased in rate. We can also not infer much information about subclasses of the phenotype *P*; we can only infer that the organism with *X* would also be annotated to any superclass of *P*. In the second case, however, we can infer that *X* would also be annotated with all subclasses of *P* (but not with its superclasses).

To avoid ambiguity in interpretation of phenotype annotations, it would be beneficial to make their intended meaning clear, in particular as the inferences that can be drawn from the interpretations are different. There have already been some efforts to integrate annotations and ontologies in a single knowledge-based model (Santana da Silva *et al.*, 2017; Hoehndorf *et al.*, 2016) which can be used as a formalized data model. Further work on formalizing the intended meaning of annotations, and the adoption of a semantic model, would further improve interoperability and reuse of these annotations.

## 5 Conclusions

We have developed a novel rule-based method for predicting phenotypes from functions. Our approach can be used as a method to validate phenotype annotations in literature-curated databases, and also to predict phenotypes from a loss of function genotype in a reverse genetics manner (Gilchrist and Haughn, 2010). While the prediction of phenotypes from genotypes is going to remain a challenge, our approach has implications for computational methods that utilize phenotypes. We demonstrated that the phenotypes we predict are predictive of interactions and of gene–disease associations; using a multi-step method in which we first predict protein functions from sequence and then phenotypes from the functions, we could predict phenotypes for genes which have not yet been investigated using a reverse genetic screen. Our approach can therefore extend the scope of phenotype-based methods, including methods for predicting variants, disease genes, or candidate drugs, to cover a significantly larger portion of the mammalian phenome.

